# Lateral prefrontal cortex as a hub for music production with gradation from structural rules to movement sequences

**DOI:** 10.1101/2020.10.21.348243

**Authors:** R. Bianco, G. Novembre, H. Ringer, N. Kohler, P.E. Keller, A. Villringer, D. Sammler

**Affiliations:** Ear Institute, University College London, UK; Max Planck Institute for Human Cognitive and Brain Sciences, Leipzig, Germany; Neuroscience and Behaviour Laboratories, Italian Institute of Technology (IIT), Rome, Italy; The MARCS Institute for Brain, Behaviour and Development, Western Sydney University, Sydney, Australia; Max Planck Institute for Empirical Aesthetics, Frankfurt am Main, Germany

**Keywords:** action hierarchy, music production, prefrontal cortex, motor sequences, abstract structure, kinematics

## Abstract

Complex sequential behaviours, such as speaking or playing music, often entail the flexible, rule-based chaining of single acts. However, it remains unclear how the brain translates abstract structural rules into concrete series of movements. Here we demonstrate a multi-level contribution of anatomically distinct cognitive and motor networks to the execution of novel musical sequences. We combined functional and diffusion-weighted neuroimaging to dissociate high-level structural and low-level motor planning of musical chord sequences executed on a piano. Fronto-temporal and fronto-parietal neural networks were involved when sequences violated pianists’ structural or motor plans, respectively. Prefrontal cortex is identified as a hub where both networks converge within an anterior-to-posterior gradient of action control linking abstract structural rules to concrete movement sequences.

## Introduction

Music, like speech, is a human-specific sequential behaviour that is governed by combinatorial structural rules. Recent research suggests that these rules not only guide music *perception* (Cheung et al. 2019), but also drive performers’ movements during music *production* (Bianco, Novembre, Keller, Kim, et al. 2016; Bianco, Novembre, Keller, Scharf, et al. 2016). It remains unclear, however, how the brain applies these rules to motor behaviour. Influential models of motor control assume that complex movement sequences are represented at different levels of abstraction – from global sequence structure to elementary movements – in topographically distinct cortical regions (Diedrichsen and Kornysheva 2015; Yokoi and Diedrichsen 2019). However, these models have so far focused on *well-learned* movement sequences, failing to account for the level of abstract rules that generate spontaneous sequential behaviour. Here, we combine models of music cognition and motor control to identify and compare the neural networks for abstract structural vs. concrete motoric representations of action sequences using a music production task.

Music is built on combinatorial structural rules, for example, the rules of harmony (Lerdahl and Jackendoff 1983; Swain 1995; Rohrmeier 2011). A musician who is able to flexibly chain tones and chords according to these rules can produce virtually infinite numbers of different musical sequences and melodies. Similar to grammatical rules in language that define *sentence* structure, these rules define which musical elements are likely to follow in a given context depending on local and non-local dependencies in an abstract *musical* structure (Patel 2003).

A wealth of research has established that listeners continuously apply these rules to the music they *hear* to derive its structure and to form expectations about what comes next (Tillmann 2012; Pearce 2018; Koelsch et al. 2019). More recently, experienced performers were found to similarly rely on these structural rules to *motorically* anticipate future elements in the music they *play*, no matter whether pieces were previously rehearsed (Palmer and van de Sande 1993, 1995), improvised (Clarke 2001), or imitated following others (Novembre and Keller 2011; Sammler et al. 2013). This indicates that playing music is not just a predetermined chain of movements, but entails several hierarchical levels of action planning: Superordinate rules regulate the combination of subordinate structural units (notes or chords), which in turn constrains the choice of different movements for execution (e.g., different finger configurations on a keyboard) (Lashley 1951; Clarke 2001; Palmer and Pfordresher 2003; Rosenbaum et al. 2007; Fitch and Martins 2014). This way, high-level musical structure can facilitate the lower-level planning of elementary movements resulting in faster and more accurate sequence execution (Palmer and Pfordresher 2003; Ahlheim et al. 2016; Bianco, Novembre, Keller, Scharf, et al. 2016).

This multi-level organization of musical action plans integrates into influential models of general action control (Fuster 2001; Cooper and Shallice 2006; Grafton and Hamilton 2007; Koechlin and Summerfield 2007; Badre and Nee 2018). These models posit anatomically distinct representations of different levels of the action hierarchy held to support action control at different time scales (Uithol et al. 2012; Hasson et al. 2015; Burt et al. 2018) and levels of abstraction (Grafton and Hamilton 2007; Badre and Nee 2018). Accordingly, enduring abstract representations of the global action structure (e.g., how to prepare coffee) incrementally activate single acts and concrete movements at shorter time scales (e.g., how to grasp the spoon to fill coffee powder into the machine) (Lashley 1951; Rosenbaum et al. 2007; Cisek and Kalaska 2010). It is well established that individual movements are represented in primary motor cortex (Yokoi et al. 2018), while higher-order representations of action sequences extend further anterior into premotor cortex (PMC) and inferior/middle frontal gyrus (IFG / MFG), as well as further posterior into inferior/superior parietal lobule (IPL / SPL) (Koechlin and Jubault 2006; Bianco, Novembre, Keller, Kim, et al. 2016; Yokoi and Diedrichsen 2019). However, the highest representational levels under investigation typically lack the flexible rule-based arrangement of single acts and movements outlined above, but remain limited to fixed combinations of motor chunks that precisely map onto well-learned sequences of finger movements (Hikosaka et al. 2002; Koechlin and Jubault 2006; Doyon 2008; Martins et al. 2019; Yokoi and Diedrichsen 2019). Exactly how the brain generates novel sequences based on abstract combinatorial rules and which neural networks underlie this ability remains largely unexplored (Ahlheim et al. 2016; Martins et al. 2019), despite the importance of flexible, non-habitual action sequences for people’s everyday life and communication (De Renzi and Lucchelli 1988; Clerget et al. 2009; Foundas and Duncan 2019).

Here, we set out to fill this gap by means of a music production task that capitalizes on the ability of musicians to rely on the structural rules of harmony when planning novel musical sequences. Although pianists develop familiarity with movement transitions between frequently co-occurring chords, structural rules do not directly map onto fixed movements. Rules allow to online anticipate the most probable next *structural* element in a given musical context (Patel 2003; Pearce 2018; Koelsch et al. 2019), for example a C major chord at the end of a C major sequence. But this chord can be implemented differently, e.g., as pressing c-e-g, e-g-c, or g-c-e keys on the keyboard, and with flexible choice of fingers for these keys. Together, these characteristics of music performance allow us to tease rule-based structural planning and motor planning apart (Clarke 2001; Palmer and Pfordresher 2003; Novembre and Keller 2011; Sammler et al. 2013; Bianco, Novembre, Keller, Scharf, et al. 2016).

We acquired behavioural, functional and diffusion-weighted neuroimaging data from expert pianists instructed to perform novel musical chord sequences on an MR-compatible piano without sound (Figure 1D). Performance was guided by series of photos of a pianist’s hand (Figure 1A-C). Sequences were constructed so as to promote planning both of (i) the next chord (i.e., which piano keys) based on the structural rules of music, and (ii) of the next movement (i.e., which fingers) based on familiar motor patterns. Violations of structural rules activated a fronto-temporal network, whilst violations of motor patterns activated a premotor-parietal network. Notably, lateral prefrontal cortex (PFC) was part of both networks and displayed an anterior-to-posterior gradient from high-level structural to low-level motor plans. PFC may thus constitute the hub at the interface between cognitive and motor networks where abstract structural rules are converted into concrete sequential movements.

**Figure1.**
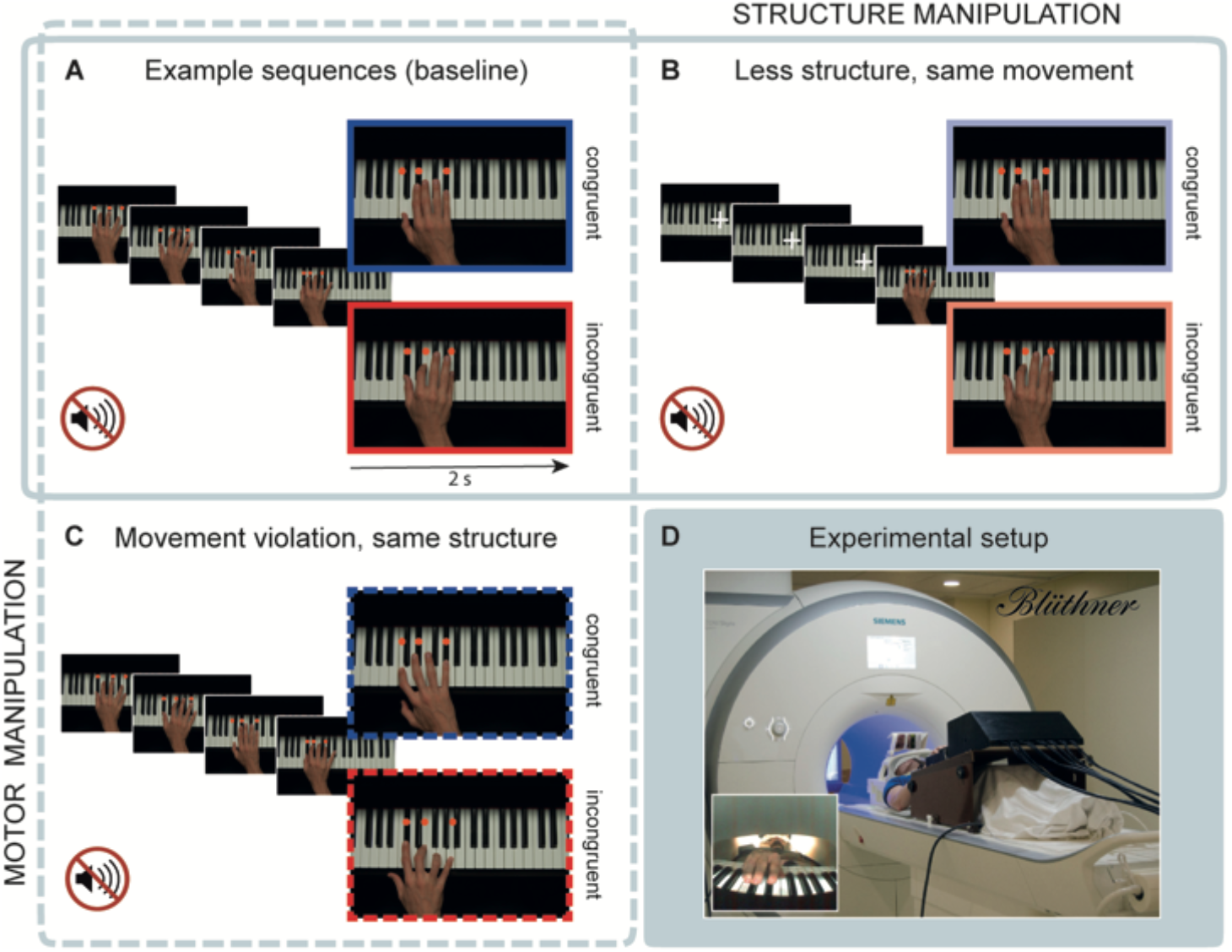
Experimental design and setting. Pianists executed novel chord sequences as shown in series of photos on screen (A-C). **(A)** The last chord of all sequences (enlarged photos in A-C) was manipulated in its structural congruency (congruent / incongruent) and was embedded in 5-chord sequences (**‘baseline’** blocks). **(B)** To address the influence of structural rules on action planning while keeping motor planning controlled, the structurally congruent and incongruent chords where embedded in a short context in **‘structure’** blocks. **(C)** To address low-level motor planning while keeping structural planning controlled, structurally congruent and incongruent chords had to be performed with an additional motor violation (i.e., with unfamiliar fingering) in **‘motor’** blocks. Effects of structure and motor manipulation were assessed in separate models (solid and dashed frame, respectively). **(D)** Pianists executed these chord sequences on an MR-compatible piano in the scanner by imitating the hand in the photos both in terms of the keys pressed and the fingering used to play. No sound was played to avoid confounding brain activity associated with the auditory processing of music. The pianist’s hand was filmed with a fish lens camera (inset on the bottom left).

## Results

We extended an established paradigm (Novembre and Keller 2011; Bianco, Novembre, Keller, Scharf, et al. 2016; Bianco et al. 2018) by adding two crucial manipulations to isolate structural and motor planning during piano performance. During the ‘baseline’ blocks of the fMRI experiment (Figure 1A), 29 pianists played unrehearsed 5-chord sequences on an MR-compatible piano by imitating, with their right-hand, actions of a model hand shown in series of photos. They were instructed to press the same keys using the same fingers as depicted. Participants played without sound to avoid confounding brain activity related to the auditory processing of music. Sequences were all different and composed so as to prime expectations about the final chord. Expectations were violated in half of the trials by introducing a structurally incongruent chord.

This manipulation has dual effects: It violates the rule-based structure of the sequence, but also disrupts frequently practiced, overlearned motor patterns between penultimate and final chords. To isolate structural planning while controlling for motor planning, the first three chords in the sequences were omitted in ‘structure’ blocks (Figure 1B) such that final chords were now preceded by only one element. This induces weaker structural expectations on the identity of the final chord than in the long sequences, while keeping the motor patterns of penultimate and final chord identical. Brain areas associated with structure processing should show stronger activity changes for incongruent chords at the end of long, as opposed to short, sequences. Second, to isolate brain regions for motor planning while keeping structure controlled, the last chord had to be played with highly uncommon finger configurations in ‘motor’ blocks (Figure 1C). These movements disrupted familiar motor patterns, that is, frequently practiced motor transitions between chords that musicians overlearn during their career (Candidi et al. 2014; Bianco, Novembre, Keller, Scharf, et al. 2016; Bianco et al. 2018). Brain areas that support motor planning should show overall stronger activity for final chords disrupting motor patterns than in the baseline sequences. Finally, to dissociate connectivity patterns of structural and motor levels of action control, we used activation peaks of the two fMRI analyses as seed and target regions in probabilistic tractography and estimated the most likely underlying white matter pathways.

### Structural planning

Behavioural data from ‘baseline and ‘structure’ blocks are shown in Figure 2A. Structurally incongruent chords were performed overall more slowly than congruent chords [main effect of STRUCTURE: *F*(1,27) = 35.79, *p* < .001, np^2^ = .57], but more so in the long than in the short context [interaction of STRUCTURE × CONTEXT: *F*(1,27) = 34.51, *p* < .001, np^2^ = .56; no main effect of CONTEXT [*F*(1,27) = 0.01, *p* = .931, np^2^ < .01]. Accuracy data showed a qualitatively similar pattern: Structurally incongruent chords were imitated less accurately than congruent chords, more so in the long than the short context, although statistics did not reach significance [interaction of STRUCTURE × CONTEXT: *F*(1,27) = 3.11, *p* = .089, np^2^ = .1; no main effect of STRUCTURE: *F*(1,27) = 2.30, *p* = .141, np^2^ = .08; no main effect of CONTEXT: *F*(1,27) = 2.55, *p* = .122, np^2^ = .09]. Overall, the interaction of structure with context suggests that structural plans were stronger in long than short sequences leading to longer reaction times (RT) and accuracy costs when the plan had to be revised in case of incongruent endings.

**Figure 2.**
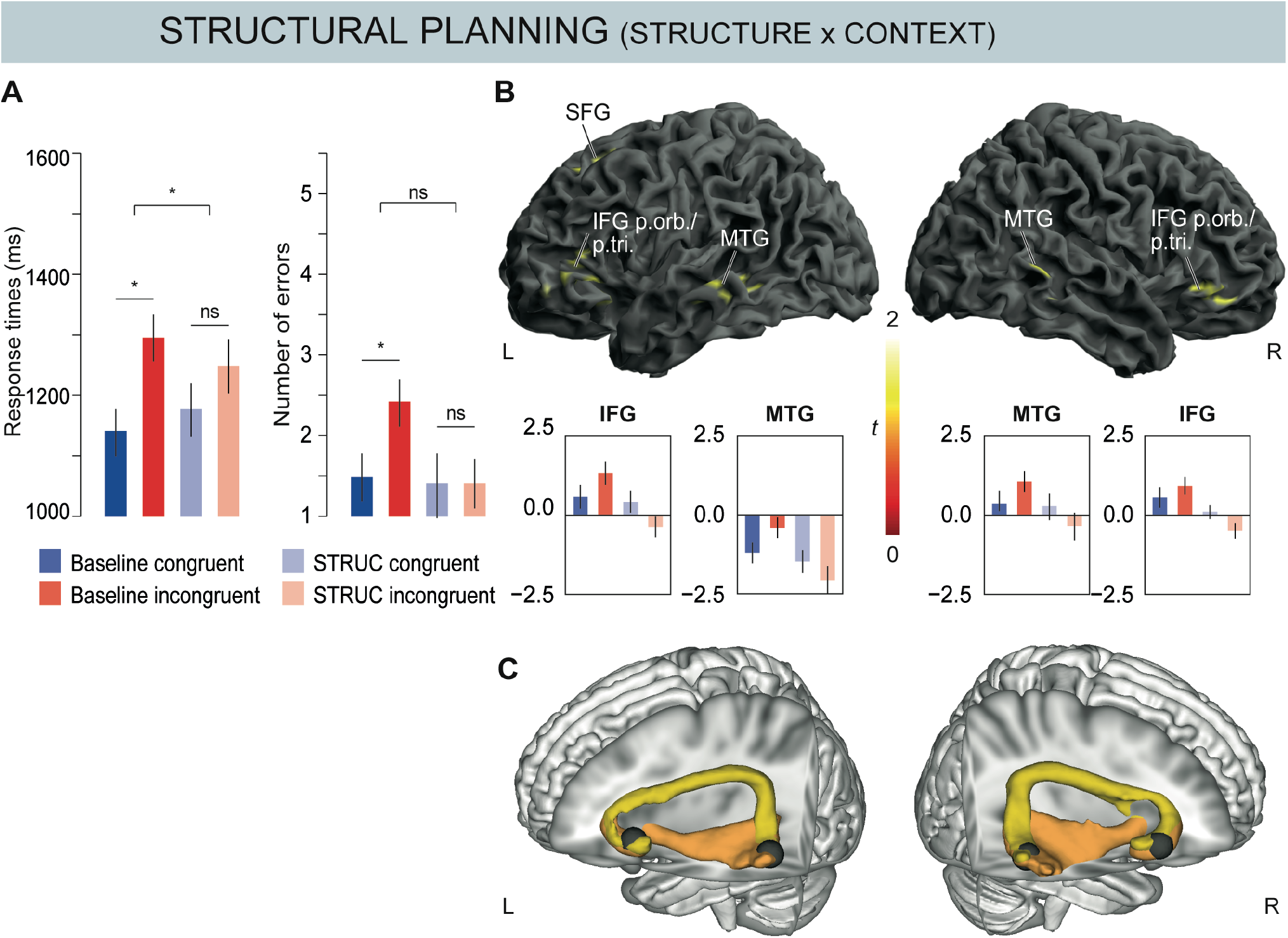
Structural planning. **(A) Behavioural performance.** Mean response times (RTs) and number of errors during execution of structurally congruent and incongruent chords in the long and short contexts (‘baseline’ vs. ‘structure’ blocks). Error bars indicate ± 1 SEM. * indicates significant effects (p < .05), ns denotes non-significant differences (p > .05). STRUC: ‘structure’ block. **(B) fMRI data.** Flexible factorial analysis of trials from the ‘baseline’ and ‘structure’ blocks with the factors STRUCTURE (congruent / incongruent) and CONTEXT (long / short). Execution of structurally incongruent chords evoked stronger activity than congruent chords when embedded in the long compared with the short context, as revealed by the interaction of STRUCTURE × CONTEXT. Activated regions included bilateral inferior frontal gyrus (IFG) and middle temporal gyrus (MTG); contrast estimates at 90% C.I. from local maxima are depicted below the brain plots. From left to right these were: left IFG (p. orb. / BA47) [−44 24 −4], left MTG (BA21) [−56 −44 −6], right MTG (BA21) [56 −36 −8], right IFG (p. orb. / BA47) [46 30 −6]. Error bars indicate ±1 SEM. Threshold for display: p_voxel_ < .001; cluster extent ≥ 46 re-sampled voxels corresponding to p_cluster_ < .05 according to Slotnick et al. (2003). p. tri.: pars triangularis; p. orb.: pars orbitalis; SFG: superior frontal gyrus. **(C) Probabilistic tractography**. Group overlay of dorsal (yellow) and ventral fibre tracts (orange) connecting anterior IFG and posterior MTG. Only voxels with fibres in more than 50% of the participants are depicted. Seed regions for probabilistic tractography are coloured in grey. Visualization of the fibre tracts was done in brainGL (http://braingl.googlecode.com). AF/SLF: arcuate / superior longitudinal fascicle; IFOF: inferior fronto-occipital fascicle.

Figure 2B shows the brain results of the flexible factorial analysis of the ‘baseline’ and ‘structure’ blocks with the factors STRUCTURE (congruent / incongruent) and CONTEXT (long / short) (see methods). We focused on the interaction of STRUCTURE × CONTEXT, that is, greater activity differences between incongruent and congruent chords in the long than in the short context because this interaction identifies brain areas involved in rule-based structural planning controlled for motor planning. We found such an interaction in bilateral anterior IFG (pars triangularis and orbitalis, BA45/47), left superior frontal gyrus (SFG), and bilateral middle temporal gyrus (MTG, BA 21) (Table 1). The activity pattern in all these clusters consistently showed higher activity for incongruent compared to congruent chords when embedded in a long context (compare red and blue bars in the parameter estimates in Figure 2B). Probabilistic fibre tractography with seeds in bilateral IFG and MTG showed that these regions are structurally interconnected both dorsally via the arcuate/superior longitudinal fascicle (AF/SLF III) as well as ventrally via the inferior fronto-occipital fascicle (IFOF; see Figure 2C).

**Table 1.** Structural planning controlled for movements (interaction of structure and context).

| Gyrus or region | Hem | BA | k | x | y | z | Z-value |
| --- | --- | --- | --- | --- | --- | --- | --- |
| **Frontal Mid.** | **L** | **46** | **387** | **-46** | **46** | **-10** | **3.98** |
| Frontal Inf. (pars Orbitalis) |  | 47 |  | -44 | 24 | -4 | 3.80 |
| Frontal Inf. (pars Triangularis) |  | 45 |  | -42 | 30 | 8 | 3.74 |
| **Frontal Inf. (pars Orbitalis)** | **R** | **47** | **109** | **46** | **40** | **-8** | **3.92** |
| " |  |  |  | 46 | 30 | -6 | 3.54 |
| **Supplementary Motor area** | **L** | **6** | **257** | **-10** | **22** | **50** | **3.91** |
| Frontal Sup. |  | 8 |  | -16 | 18 | 54 | 3.90 |
| Frontal Sup. Medial |  | 8/32 |  | -8 | 30 | 46 | 3.45 |
| **Temporal Mid.** | **L** | **21** | **108** | **-56** | **-44** | **-6** | **3.85** |
| Temporal Inf. |  | 20 |  | -52 | -50 | -14 | 3.64 |
| Temporal Mid. |  | 21 |  | -60 | -36 | -10 | 3.45 |
| **Temporal Sup.** | **L** | **22** | **89** | **-60** | **-22** | **-6** | **3.83** |
| **Temporal Mid.** | **R** | **21** | **88** | **56** | **-36** | **-8** | **3.57** |

Whole-brain activation cluster sizes (*k*), MNI coordinates (*x*, *y*, *z*), and Z-scores for the Structure x Context contrast ( $p_{\text{voxel}} < .001$ ; correction for multiple comparisons to $p < .05$ was obtained using a voxel cluster extent threshold procedure which led to minimum cluster extent threshold of 46 re-sampled voxels). BA: Brodmann area, Hem.: hemisphere. Inf.: Inferior, Mid.: Middle.

### Motor planning

Figure 3A shows participants’ behavioural performance in ‘baseline’ and ‘motor’ blocks. Motorically unfamiliar chords were imitated more slowly and less accurately than motorically familiar chords [main effect of MOVEMENT for RTs: *F*(1,27) = 181.30, *p* < .001, np^2^ = .87; accuracy: *F*(1,27) = 20.55, *p* < .001, np^2^ = .43]. RTs were also slower and accuracy lower for structurally incongruent than congruent chords [main effect of STRUCTURE for RTs: *F*(1,27) = 36.20, *p* < .001, np^2^ = .57; accuracy: *F*(1,27) = 31.09, *p* < .001, np^2^ = .54]. Finally, we found significant interactions of STRUCTURE × MOVEMENT for RTs [*F*(1,27) = 36.72, *p* < .001, np^2^ = .58] and accuracy [*F*(1,27) = 4.58, *p* = .041, np^2^ = .15]: RTs were slower in structurally incongruent than congruent chords when movements were familiar [*t*(27) = −7.521, *p* < .001] while performance was similarly slow when movements were unfamiliar [*t*(27) = −1.167, *p* = .507]. Structurally incongruent chords induced more errors than congruent chords when movements were familiar [*t*(27) = −2.414, *p* = .005], but more so when movements were unfamiliar [*t*(27) = −5.643, *p* < .001]. This indicates that structural violations, but particularly unfamiliar fingerings, place higher demands on motor planning.

**Figure 3.**
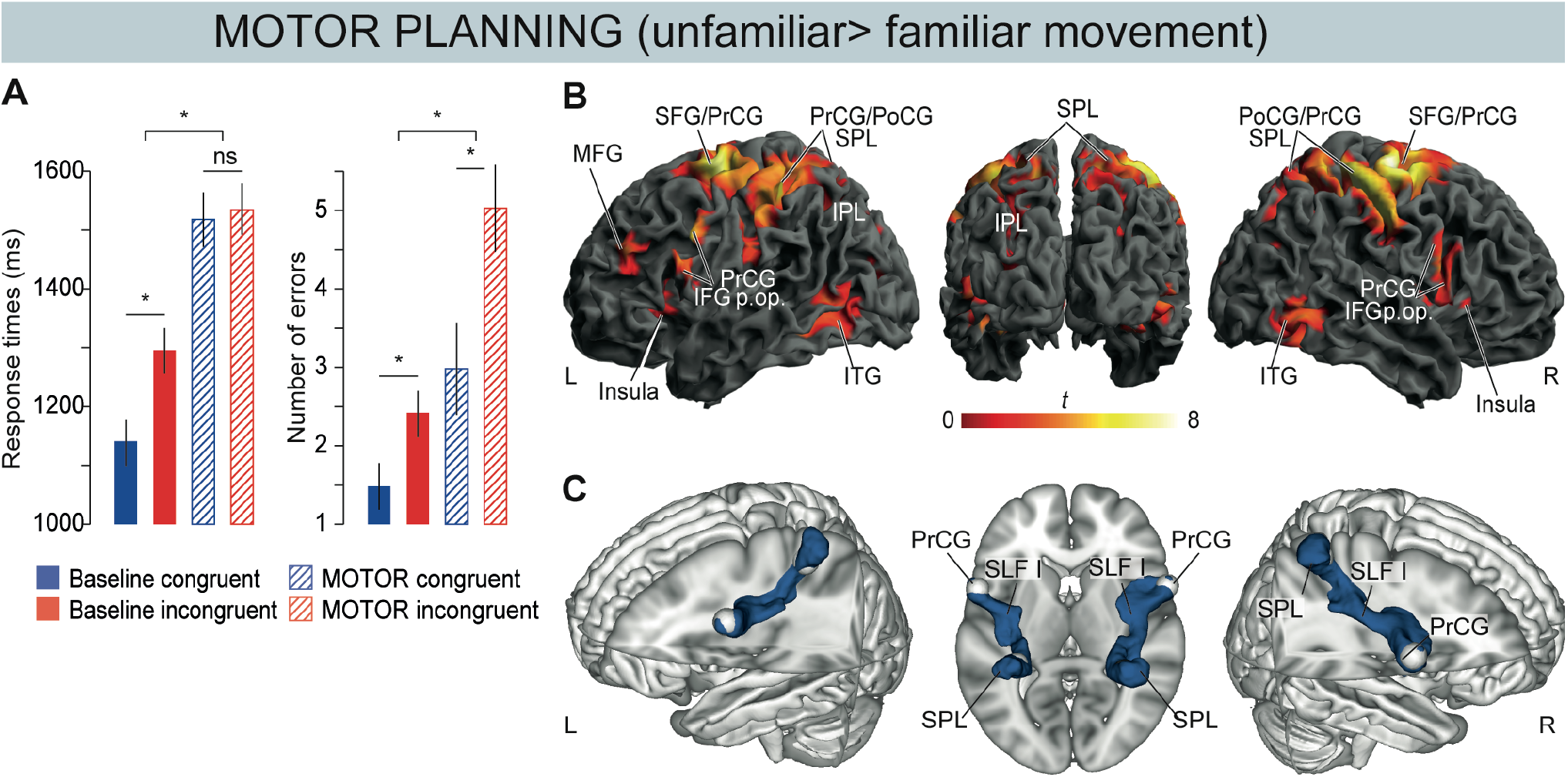
Motor planning. **(A) Behavioural performance.** Mean RTs and number of errors during motorically familiar (filled bars) or unfamiliar execution (striped bars) of structurally congruent and incongruent chords (‘baseline’ vs. ‘motor’ blocks). Error bars indicate ± 1 SEM. * indicates significant effects (p < .05), ns denotes non-significant differences (p > .05). **(B) fMRI data.** Flexible factorial analysis of the ‘baseline’ and ‘motor’ blocks with the factors STRUCTURE and MOVEMENT. Execution of chords with unfamiliar compared with familiar movements elicited stronger activity in bilateral parieto-frontal action areas. Threshold for display: p_voxel_ < .001; cluster extent ≥ 46 re-sampled voxels corresponding to p_cluster_ < .05 according to Slotnick et al. (2003). IFG: inferior frontal gyrus; p. op.: pars opercularis; MFG: middle frontal gyrus; PrCG: precentral gyrus; PoCG: postcentral gyrus; SFG: superior frontal gyrus; SPL: superior parietal lobule; IPL: inferior parietal lobule; ITG: inferior temporal gyrus. **(C) Probabilistic tractography.** Group overlay of dorsal fibre tracts (blue) connecting PrCG and SPL. Only voxels with fibres in more than 50% of the participants are depicted. Seed regions for probabilistic tractography are coloured in light grey. Visualization of the fibre tracts was done in brainGL (http://braingl.googlecode.com). SLF: superior longitudinal fascicle.

Figure 3B shows the brain results of the flexible factorial analysis of the ‘baseline’ and ‘motor’ blocks with the factors STRUCTURE and MOVEMENT. We focused on the main effect of MOVEMENT, that is, stronger activity for motorically unfamiliar than familiar finger configurations, because it identifies brain areas taxed by motor demands, in chord sequences with the same structure. We found such a main effect in a broadly distributed set of fronto-parietal regions. These included bilateral IFG (pars opercularis, BA44) and precentral cortices (PrCG, BA6), insula, SFG, and left middle frontal gyrus (MFG, BA46), as well as bilateral postcentral gyrus (PoCG), superior parietal lobule (SPL), left inferior parietal lobule (IPL), bilateral inferior temporal gyrus (ITG), and lobules VII and VIII of the cerebellum (see Table 2 and Figure 3B). Fibre tractography with seeds in bilateral PrCG and SPL showed exclusively dorsal connections via the superior longitudinal fascicle (SLF I; see Figure 3C).

**Table 2.** Motor planning controlled for structure (main effect of movement).

| Gyrus or region | Hem | BA | k | x | y | z | Z-value |
| --- | --- | --- | --- | --- | --- | --- | --- |
| Precentral | R | 6 | 16579 | **36** | **-14** | **66** | inf |
| Postcentral |  | 3b |  | 40 | -32 | 56 | 7.15 |
| Superior Parietal |  | 2 |  | 34 | -42 | 60 | 6.58 |
| Supp. Motor Area |  | 6 |  | 6 | -2 | 52 | 6.45 |
| Frontal Sup. | L | 6 |  | -22 | -6 | 64 | 6.85 |
| Supp. Motor Area |  | 6 |  | -6 | -2 | 52 | 6.36 |
| Precentral |  | 6 |  | -36 | -10 | 60 | 5.66 |
| Postcentral |  | 3 |  | -46 | -30 | 52 | 5.65 |
| **Cerebellum** | **L** | **VI** | **1495** | **-32** | **-54** | **-22** | **5.83** |
| Temporal Inf. |  | 37 |  | -46 | -50 | -14 | 5.01 |
| " |  |  |  | -44 | -62 | -2 | 4.59 |
| **Precentral/Frontal (Opercularis)** | **L** | **6/44** | **1305** | **-56** | **8** | **30** | **5.82** |
| " |  |  |  | -58 | 8 | 22 | 5.78 |
| Frontal Mid. |  | 46 |  | -44 | 40 | 30 | 5.38 |
| **Precentral/Frontal (Opercularis)** | **R** | **6/44** | **830** | **60** | **10** | **16** | **5.16** |
| Insula |  | 48 |  | 42 | 0 | 10 | 4.21 |
| " |  |  |  | 34 | 18 | 4 | 3.74 |
| **Cerebellum** | **R** | **VI** | **1108** | **30** | **-50** | **-24** | **4.94** |
| Temporal Inf. |  | 37 |  | 54 | -60 | -6 | 4.92 |
| Occipital Inf. |  | 19/37 |  | 42 | -64 | -14 | 3.74 |
| **Insula** | **L** | **48** | **510** | **-38** | **14** | **2** | **4.34** |
| " |  |  |  | -32 | 22 | -2 | 4.25 |
| " |  |  |  | -32 | 18 | 8 | 4.08 |
| **Cerebellum** | **L** | **VIIb** | **88** | **-6** | **-72** | **-42** | **4.04** |
| " |  | VIIIa |  | -14 | -66 | -48 | 4.09 |
| " |  | VIIb |  | -14 | -74 | -46 | 3.61 |
| **Occipital Mid.** | **L** | **19** | **141** | **-32** | **-80** | **10** | **3.97** |
| " |  |  |  | -32 | -82 | 24 | 3.27 |

Whole-brain activation cluster sizes ( $k$ ), MNI coordinates ( $x, y, z$ ), and Z-scores for the Unfamiliar > Familiar Movement contrast ( $p_{\text{voxel}} < .001$ ; correction for multiple comparisons to $p < .05$ was obtained using a voxel cluster extent threshold procedure which led to minimum cluster extent threshold of 46 re-sampled voxels). BA: Brodmann area, Hem.: hemisphere. Inf.: Inferior, Mid.: Middle, Sp.: Superior.

### Dissociation of structural vs. motor levels of action planning

Figure 4 (central panels) summarizes our findings showing the fronto-temporal network for structural planning (yellow) and the fronto-parietal network for motor planning (blue) of musical actions. The two networks hardly overlap. Zooming in on frontal regions (outer panels) further illustrates more fine-grained dissociations along the anterior-to-posterior axis of lateral prefrontal cortex: rule-based structural planning was best captured by activity in anterior IFG (pars triangularis and orbitalis, BA 45/BA47), whilst motor planning evoked bilateral activity in posterior IFG (pars opercularis, BA 44) and premotor cortices (PrCG, BA 6).

**Figure 4.**
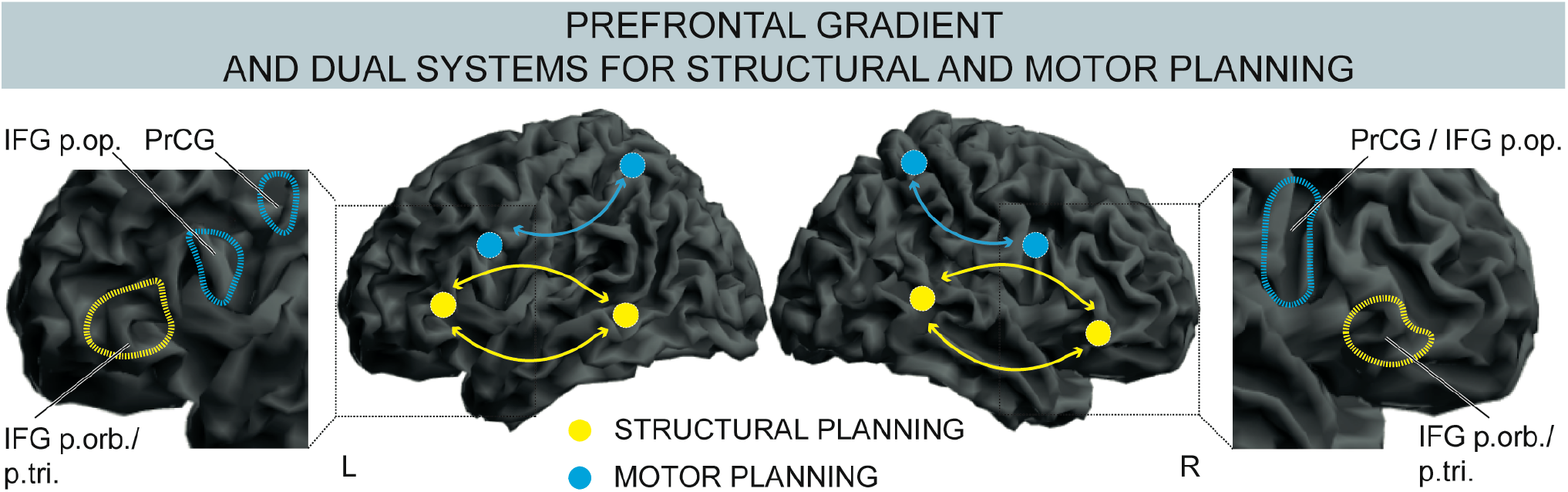
Dual networks and prefrontal gradient for structural and motor plans during music production. Anterior frontal and posterior temporal regions connected via dorsal (AF/SLF III) and ventral pathways (IFOF) support rule-based structure processing during production of novel musical sequences (yellow). Conversely, posterior frontal and parietal regions dorsally connected via SLF I support the planning and execution of motor movements. The insets on the left and right suggest an anterior-to-posterior gradient along lateral prefrontal cortex from abstract structural planning in bilateral pars triangularis and orbitalis of IFG (BA45/BA47) to concrete motor planning in bilateral PrCG (BA6) and pars opercularis of IFG (BA44). IFG: inferior frontal gyrus; p. op.: pars opercularis; p. tri.: pars triangularis; p. orb.: pars orbitalis; PrCG: precentral gyrus; BA: Brodmann area.

## Discussion

We isolated the neural networks of abstract structural and concrete motor planning during music production. Pianists executed unrehearsed chord sequences on an MR-compatible piano in the scanner. The final chords of the sequences were either structurally congruent or incongruent. To isolate high-level structural planning of the ongoing action from low-level motor planning, the final target chords were embedded in shorter sequences (i.e., with less structure), and brain activity changes for incongruent chords were compared between long and short sequences. To isolate lower-level motor planning from structural planning, the final chords of sequences with identical structure had to be played with familiar or unfamiliar movements. We found two networks selectively involved in one or the other level of action planning: bilateral anterior IFG (BA45/47) and posterior MTG (BA21) were activated by structure violations in long more than in short sequences, and were interconnected both ventrally and dorsally via IFOF and AF/SLF. Bilateral posterior IFG / PrCG (BA44/6) and superior parietal areas were activated by motor violations, and were interconnected dorsally via SLF. The combined data demonstrate the multi-level contribution of anatomically distinct cognitive and motor networks to the execution of novel movement sequences, adding a rule-based level of abstraction to models of hierarchical motor control. Importantly, PFC is identified as a hub where both networks converge, and where abstract structural representations may gradually be transformed into sequential motor behaviour.

### Rule-based structural planning of sequences involves fronto-temporal network

Execution was slower for structurally incongruent than congruent chords more so in long than short sequences (Figure 2A). This indicates that structural violations were more unexpected after long contexts, implying that pianists had derived the abstract structure of the ongoing sequence using their rule knowledge, allowing them to plan structural elements ahead of time: This facilitated execution when plans were met, but induced RT costs when plans were violated and had to be revised (Novembre and Keller 2011; Sammler et al. 2013; Bianco, Novembre, Keller, Scharf, et al. 2016).

In line with the behavioural results, structural violations activated bilateral anterior IFG (BA45/47) and MTG (BA21) more strongly in long than in short sequences (Figure 2B). These findings are consistent with previous work implicating IFG together with temporal areas in the processing of structural violations during music *perception* (Zatorre et al. 1998; Maess et al. 2001; Koelsch et al. 2002, 2005; Tillmann et al. 2003, 2006; Seung et al. 2005; Vuust et al. 2011; Kim and Sikos 2011; Koelsch 2011; Sammler et al. 2011; Zatorre and Salimpoor 2013; Musso et al. 2015; Bianco, Novembre, Keller, Kim, et al. 2016; Di Liberto et al. 2020). Here, we show for the first time a similar network during music *production*. This network extended slightly more anteriorly into pars orbitalis of IFG and more ventrally into MTG compared with auditory paradigms. There, activation peaks typically fall into pars opercularis and triangularis (BA44/45) of IFG and superior temporal gyrus (STG) (Koelsch 2005), except for participants with high musical expertise who showed MTG activity, possibly reflecting their stronger music-structural representations (Zatorre et al. 1998; Janata et al. 2002; Haslinger et al. 2005; Seung et al. 2005; Bianco, Novembre, Keller, Kim, et al. 2016). We consider it unlikely that the present anterior shift in PFC (and ventral shift in the temporal lobe) is a matter of modality (perception vs. production). Rather we attribute it to the tighter control of the structural violations for low-level (sensory and motor) features in the present than in previous fMRI studies, by comparing effects between long and short sequences. This may have better dissociated abstract structural from (spurious) sensorimotor processes (see also below).

Diffusion data revealed both the arcuate / superior longitudinal fascicle (AF/SFL) and the inferior fronto-occipital fascicle (IFOF; Figure 2C) as likely network connections between IFG and MTG. In music perception, both these pathways have been associated with the ability to process musical structure (Musso et al. 2015), which is impaired after damage to these fibre tracts in acquired and congenital amusia (AF/SLF: Loui et al. 2009; Peretz 2016; Sihvonen et al. 2017, but see Chen et al. 2015; IFOF: Sihvonen et al. 2017). In music production, available tractography studies have focused mostly on audio-motor coupling (Halwani et al. 2011; Engel et al. 2014) rather than structural processing. Hence, the present data are the first to highlight bilateral ventral and dorsal pathways linking IFG and MTG as most likely anatomical scaffold to support knowledge-driven, rule-based computations not only in perception but also in production of novel musical sequences.

### Motor planning involves a fronto-parietal network

Performance was overall slower when chords had to be executed with uncommon fingerings (Figure 3A), reflecting higher demands on motor planning when familiar movement transitions were undue. The brain data mirrored these results in a fronto-parietal network including posterior IFG and PrCG (BA44/6), primary motor areas and the SPL / IPL (Figure 3B). Those were interconnected via the most dorsal branch of the SLF (Figure 3C), crucial for hand motor control and movement selection (Schulz et al. 2015). This network has been previously associated with motor preparation, prediction of motor outcomes or integration of sensorimotor information and has been found in studies investigating violations of simple actions (e.g., a whole-hand grip for picking up a pin) (Grèzes and Decety 2001; Williams et al. 2007; Papitto et al. 2020), or the neural representation of overlearned sequential finger movements (Yokoi et al. 2018; Yokoi and Diedrichsen 2019). In particular, Yokoi et al. (2018; 2019) demonstrated that fixed, overlearned chunks of familiar sequential finger movements were represented in parietal (including SPL / IPL) and posterior frontal regions including BA6 and BA44 (see also (Farag et al. 2010)), similar to the regions we found activated when familiar, overlearned movement transitions were disrupted.

Interestingly, the present results also resemble our previous findings of functionally coupled posterior IFG and SPL during execution of *structurally* incongruent chords, despite standard movement transitions (i.e., corresponding to the simple contrast of incongruent minus congruent ‘baseline’ sequences, as in Figure S1) (Bianco, Novembre, Keller, Kim, et al. 2016). This suggests that frequently co-occurring structural elements can consolidate in overlearned motor patterns, i.e., fixed chunks that are concurrently disrupted in case of a structural violation. Indeed, motor training has been shown to shift processing from higher abstract to lower motoric levels of the action hierarchy facilitating sequence execution (Diedrichsen and Kornysheva 2015). This further highlights the importance of the short sequences in our paradigm for dissociating fronto-temporal structure-related from fronto-parietal movement-related representations of action sequences, allowing to uncover an additional abstract and cognitive level of action planning that guides the flexible, rule-based execution of novel movement sequences.

### Anterior-to-posterior frontal gradient: from structural rules to movement sequences

The networks associated with high-level structure and low-level motor planning involved anterior and posterior prefrontal regions, respectively. This observation is reminiscent of functional anterior-to-posterior gradients in PFC postulated by models of action control (Badre and Nee 2018; Rouault and Koechlin 2018). Action control is generally referred to as the ability to select and coordinate actions or thoughts in relation to internal higher-level goals, and is a cardinal function of the lateral PFC (Miller and Cohen 2001; Koechlin et al. 2003; Wood and Grafman 2003). Neuroimaging studies revealed progressively more anterior activity in lateral PFC during response selection to progressively more complex stimuli (reviewed by Koechlin & Summerfield, 2007). For example, while movement sequences in response to simple sensory cues evoked activity in motor and premotor regions (BA4/6) (Koechlin et al. 2003; Koechlin and Jubault 2006; Badre and D’Esposito 2007), more complex sequences coordinated across nested temporal frames (Nee and D’Esposito 2017) or embedded in hierarchical patterns (Koechlin and Jubault 2006) recruited anterior frontal regions (BA44/45/9 up to BA46/10). Our anterior and posterior IFG clusters are consistent with these findings and suggest that IFG may constitute the hub where conceptual and motor networks converge, and abstract structural representations are gradually transformed into sequential motor behaviour. Tracking the temporal and causal dynamics of this assumed transformation along the gradient and over the course of learning are interesting prospects for future research.

### Parallels to speech production

The idea that IFG is a key region at the interface between cognitive and motor networks finds noteworthy parallels in recent models of speech production (Garagnani and Pulvermüller 2013; Flinker and Knight 2018; Hage 2018; Matchin and Hickok 2019). They suggest that the conversion of abstract linguistic structures, such as sentences or words, into chains of articulatory movements relies on the interaction between the MTG as a syntactic hub (Meyer and Friederici 2016), and anterior frontal regions (in particular pars triangularis of IFG) as the interface to motor implementations of speech in premotor and motor areas. These models are supported by studies showing abstract linguistic information processing in anterior frontal regions (syntactic sentence structure in BA45: Segaert et al. 2013; word generation and syllabification in BA44: Indefrey and Levelt 2004; Bourguignon 2014; Flinker et al. 2015) and lower-level articulatory programs represented in precentral areas (Flinker et al., 2015; see also Uddén & Bahlmann, 2012). Moreover, IFG activity was found to be more posterior in case of highly automatic linguistic structure processing in one’s first compared with one’s second language (Jeon and Friederici 2015). This resonates with the above-raised idea that frequently co-occurring structural elements may consolidate in overlearned chunks that are represented more posteriorly in the frontal hierarchy. Overall, these parallels between speech and music production are in line with ideas that the IFG plays a domain-general role in controlling sequential behaviours (Fitch and Martins 2014; Bornkessel-Schlesewsky et al. 2015; Rouault and Koechlin 2018) by acting as a multimodal association zone or “cortical hub” (Friederici and Singer 2015) that links cognitive and sensorimotor networks and thus supports the formation of distributed action circuits.

## Conclusion

Understanding how actions are neurally represented at their different hierarchical levels is a first crucial step to understand what enables us to flexibly generate the variety of action sequences we use every day to communicate and interact with the world. Whilst exciting theoretical and experimental advances have been achieved in understanding the generative power of human-specific, rule-based cognitive abilities (Wilson et al., 2017), it remains an open question why we ultimately select one action sequence, and not another, amongst the infinite range of sequences we are able to conceive. Our work with musical sequences grounds the anatomical bases of production of rule-based actions within a dual network architecture. Further studies are needed to detail the timing of the recruitment of these networks, and how the constituent brain areas interact with one another during music production. Research in this direction can increase our understanding of how complex sequential behaviours like speech and music gain the flexibility needed for meeting the demands of real-life interaction.

## Methods

### Participants

We present data from 29 pianists (10 female; mean age = 25.5 years, *SD* = 3.9). They had a minimum of 5 years of piano training in classical Western tonal music (range = 6-28 years, mean = 17.0 years, *SD* = 5.2) and had started to play the piano at an average age of 8.5 years (*SD* = 3.4, range = 5-17 years). Data were acquired from eight more pianists that were however excluded from the analysis because they were not able to perform the task or because of technical problems during data acquisition. Written informed consent was obtained from each participant prior to the study that was approved by the local ethics committee of the University of Leipzig (016-15-26012015).

### Stimuli

Stimuli consisted of 26 different 5-chord sequences that were presented as photos of a hand playing chords on a piano (i.e., one photo for each chord; Figure 1; see also Bianco et al., 2016, 2018). Each chord was presented with the same duration of 2 s. The interval between sequences was jittered between 3 and 9 s (mean = 5.6 s) during which a black screen was displayed. The sequences were composed of five chords according to the rules of classical harmony in six different tonalities (D, E, Bb, Ab, A and Eb major). Each chord consisted of three notes to be played with the right hand.

We presented three variants of these chord sequences in separate blocks. First, ‘baseline’ blocks (Figure 1A) corresponded to the paradigm of Bianco et al. (2016), that is, 5-chord sequences that ended with either a structurally congruent (a Tonic chord) or an incongruent chord (a Neapolitan chord, namely a minor subdominant with a diminished sixth instead of a fifth, rarely used in classical harmony to resolve a musical sequence). Final chords were controlled for visual appearance as in Bianco et al. (2016) by balancing the average amount of black and white keys in congruent and incongruent chords. Furthermore, chords that appeared as congruent Tonic in one sequence also appeared as incongruent Neapolitan in another sequence. However, given any chord preceding the final one, motor plans and movement trajectories towards congruent and incongruent final chords inevitably differ, conflating motor and abstract structural planning.

To isolate brain regions involved in abstract structural planning while controlling for these motoric differences, 5-chord sequences were truncated and only the last two chords of each sequence were presented in ‘structure’ blocks (Figure 1B: structure manipulation). This way, any motoric differences between structurally congruent and incongruent chords were identical in long and short sequences that, hence, differed only in the strength of the structure plan. Consequently, brain regions showing stronger activity differences between incongruent and congruent chords in long than short sequences must be linked to abstract structural planning. To avoid general differences in motor activity between long and short sequences due to a different number of movements preceding the final target chords, the two photos constituting the short sequences were preceded by three photos of an empty piano and a white fixation cross: During the presentation of these, participants had to perform a thumb opposition task which was monitored online through an MR-compatible camera.

Finally, to isolate brain regions involved in low-level motor planning while controlling for abstract structural planning, the final chords of the long sequences had to be played with unusual finger configurations in ‘motor’ blocks (Figure 1C: motor manipulation). Such finger configurations were anatomically awkward and highly unlikely to be used, as revealed by expert pianists’ rating on their unconventionality in Bianco et al., 2016. Note that the context and, hence, abstract structural planning in ‘motor’ blocks was the same as in ‘baseline’ blocks. In other words, these blocks differed only in movement demands such that their direct comparison should isolate brain areas associated with more effortful motor planning.

### Procedure

Pianists were instructed to watch and simultaneously imitate the chord sequences played by the hand in the photos: They were instructed to reproduce both, keys pressed and fingering used. A mirror mounted on the head coil allowed them to see the photos projected onto a screen at the head-end of the MR-scanner. For execution, they used a custom-built MR-compatible piano with 27 weighted keys manufactured by Julius Blüthner Pianofortefabrik GmbH (Leipzig, Germany; Figure 1D). Weighted keys increase the ecological validity of the performance by giving the users a similar touch experience as playing on a real piano. The experiment was run in the absence of musical sound, that is, participants played the chord sequences without receiving auditory feedback of their motor actions (and likewise, no sounds were associated with the photos). Key presses, velocity, and key releases were sensed optically using a light-emitting diode, a matching phototransistor, a pair of fiber optic cables, and a reflector for each key of the MR-piano as in (Hollinger et al. 2007). All electronic components of the piano were located in the room adjacent to the scanning room, with the optical cables entering the scanning room through the wall (waveguide). The piano was positioned on a slightly tilted wooden stand over the participant lying supine in the bore of the MR scanner. Pianists’ finger movements were monitored and recorded through an MR-compatible camera with fisheye lens (12M camera, MRC Systems, Heidelberg, Germany) placed on top of the piano. This allowed offline analysis of fingering errors committed by pianists.

The 26 stimuli were repeated in all their manipulation variants across 6 blocks. All blocks (each approximately 8 minutes) consisted of 26 trials and were organized as follows: Two blocks of the type ‘baseline’ contained long sequences with structurally congruent / incongruent final chords played with familiar movements (Figure 1A); two blocks of the type ‘structure’ contained short sequences with structurally congruent / incongruent endings also played with familiar movements (Figure 1B); in the remaining two blocks of the type ‘motor’, structurally congruent / incongruent chords at the end of long sequences were played with unfamiliar movements (Figure 1C). The order of blocks was randomized across participants. Trials within each block were presented in pseudo-random order with the constraint that no more than three sequences of the same condition followed each other. Stimulus presentation was controlled with Presentation software (version 14.9, Neurobehavioural Systems, Inc.). Pianists’ key presses on the MR-piano were recorded by custom-written Python software running on a Linux computer.

To acquaint participants with the task in the scanner, a mock training session was run about one week before the scanning day. During this pre-session, participants were trained with a different set of sequences in different tonalities (G, B, F, Db) in a mock scanner on a midi keyboard (M-Audio Keystation 49e, inMusic GmbH, Ratingen, Germany).

### MR data acquisition

The experiment was carried out in a 3.0-Tesla Siemens PRISMA whole body magnetic resonance scanner (Siemens AG, Erlangen, Germany) using a 32-radiofrequency-channel head coil. *Functional magnetic resonance images* were acquired using a T2*-weighted 2D echo planar imaging (EPI) sequence with TE = 30 ms and TR = 2000 ms. 240 volumes were acquired for each block, with a square FOV of 210 mm, with 37 interleaved slices of 3.2 mm thickness and 15% gap (3 × 3 × 3.68 mm^3^ voxel size) aligned to the AC-PC plane, and a flip angle of 77°.

High-resolution *T1-weighted anatomical images* and *diffusion-weighted images* of the participants were either taken from the database of the Max Planck Institute or acquired in the context of the fMRI experiment. Diffusion-weighted MR data were available for 27 pianists. Anatomical images were recorded using a 3D MP2RAGE sequence (TI_1_ = 700 ms, TI_2_ = 2500 ms, TE = 2.03 ms, TR = 5000 ms) with a matrix size of 240 × 256 × 176, with 1 mm isotropic voxel size, flip angle_1_ of 4°, flip angle_2_ of 8°, and GRAPPA acceleration factor of 3. Diffusion-weighted data were acquired with a twice-refocused spin echo EPI sequence (TE = 100 ms, TR = 12900 ms, 88 axial slices without gap, FOV = 220 mm, matrix size = 128 × 128, iPAT = 2) with 1.71875 mm isotropic voxel size. Diffusion-weighting was isotropically distributed along 60 diffusion-encoding gradient directions with a b-value of 1000 s/mm^2^. Additionally, seven images without diffusion-weighting (b0) were recorded evenly distributed across scan time and served as anatomical reference for offline motion correction.

### Behavioural data analysis

Performance of the last chord was analysed as in previous studies using this paradigm (Novembre and Keller 2011; Sammler et al. 2013; Bianco, Novembre, Keller, Scharf, et al. 2016). Trials were included in the analysis when (1) the penultimate and final chord of the sequence were imitated correctly, both in terms of keys and fingering, (2) when the three keys in the penultimate and in the final chord were pressed synchronously (i.e., no more than 150 ms elapsed between the first and the last of the 3 keystrokes), and (3) when response times (RTs) of the final chord were below 3000 ms. RTs were the averages of the three keystrokes of the final chord time-locked to the onset of the last photo in the sequence. Fingering of the participants was analysed through off-line inspection of the video recordings of their hands. For each participant, RTs that deviated by more than 2 *SD*s from the mean across conditions were discarded from the analysis. Based on these exclusion criteria, an average of 69 ± *SD* 15.5 % of the total number of trials remained to be analysed per participant. RTs and number of errors (pooled key and fingering errors) were used as dependent variables.

To address the rule-based structural planning while keeping motor planning controlled, we ran an ANOVA with the repeated-measures factors STRUCTURE (congruent / incongruent) and CONTEXT (long / short) including the trials from the ‘baseline’ and ‘structure’ blocks. To address low-level motor planning while keeping structural planning controlled, we ran a two-way analysis of variance (ANOVA) with the repeated-measures factors STRUCTURE (congruent / incongruent) and MOVEMENT (familiar / unfamiliar) including the trials from the ‘baseline’ and ‘motor’ blocks. ANOVAs were implemented in the R environment (version 0.99.320) using the ‘ezANOVA’ function (Michael Lawrence, 2016). Post-hoc *t*-tests were used to resolve significant interactions, and Bonferroni-correction was applied based on the number of comparisons.

### fMRI data analysis

fMRI data were analysed with SPM12 (Welcome Trust Centre for Neuroimaging, University College, London, UK, http://www.fil.ion.ucl.ac.uk/spm/software/spm12) using standard spatial pre-processing procedures. These consisted of slice time correction (using cubic spline interpolation), spatial realignment, co-registration of functional and anatomical data (uniform tissue-contrast image masked with the 2^nd^ inversion image from the MP2RAGE sequence), spatial normalization into the MNI (Montreal Neurological Institute) stereotactic space, that included resampling to 2 × 2 × 2 mm voxel size. Finally, data were spatially low-pass filtered using a 3D Gaussian kernel with full-width at half-maximum (FWHM) of 8 mm and temporally high-pass filtered with a cut-off of 1/128 Hz to eliminate low-frequency drifts.

The evoked hemodynamic response to the onset of the final chord was modelled for each of the 6 conditions (the congruent / incongruent chords in the ‘baseline’, ‘structure’, and ‘motor’ blocks) as boxcars convolved with a hemodynamic response function (HRF). All trials were included in the brain data analysis to secure statistical power. Estimated motion re-alignment parameters were added to this design as covariates of no interest to regress out residual motion artifacts and to increase statistical sensitivity.

Whole-brain random-effects models were implemented to account for within-subject variance. Statistical parametric maps for each of the six conditions (one-sample *t*-tests against implicit baseline) were generated for each participant in the context of the general linear model (GLM) for use in the second level group analysis.

We then ran two models with 2 × 2 within-subject flexible factorial designs to dissociate brain regions associated with the different levels of the action hierarchy. The first model contained the trials from the ‘baseline’ and ‘structure’ blocks and the factors STRUCTURE (congruent / incongruent) and CONTEXT (long / short). The interaction of STRUCTURE × CONTEXT should unveil brain areas modulated by the strength of the structure plan. These regions should show stronger activity to structurally incongruent than congruent chords in the long compared with the short context. The second model with STRUCTURE and MOVEMENT as factors included the trials from the ‘baseline’ and ‘motor’ blocks. The main effect of MOVEMENT (unfamiliar > familiar) should identify brain regions involved in low-level motor plans regardless of structural congruency.

For statistical thresholding, we ran a Monte Carlo simulation in Matlab (1000 iterations, no volume mask) that suggested a cluster extent threshold of ≥ 46 resampled voxels at a voxel-level uncorrected *p*-value of .001 to ensure whole-volume type I error probability smaller than .05 (Slotnick et al., 2003; code available at http://www2.bc.edu/sd-slotnick/scripts.html). Anatomical labelling was based on the SPM anatomy toolbox (Eickhoff et al. 2005).

### Diffusion data analysis

Processing of diffusion data and anatomical reference images was done in FSL (version 5.0.9, FMRIB, University of Oxford, www.fsl.fmrib.ox.ac.uk/fsl), SPM12 and LIPSIA (Max Planck Institute for Human Cognitive and Brain Sciences, Leipzig, Germany; Lohmann et al., 2001). Diffusion-weighted images were first motion-corrected using rigid-body transformations based on the seven (b0) non-diffusion-weighted reference images, and then registered to the T1-weighted anatomical images resampled to diffusion space with 1.72 × 1.72 × 1.72 mm resolution. Subsequently, fibre orientation was estimated in each voxel by means of the software module BEDPOSTX (with standard options) in FSL using a crossing fibre model with up to two directions per voxel (Behrens, Berg, Jbabdi, Rushworth, & Woolrich, 2007).

Seed regions for tractography were obtained by first projecting MNI group coordinates in bilateral IFG [−44, 24, −4; 46, 30, −6] and MTG [−56, −44, −6; 56, −36, −8], PrCG [−56, 8, 30; 60, 10, 16] and SPL [−32, −38, 54; 34, −42, 60] into each participants’ diffusion MRI space. Coordinates that fell into sulci or grey matter were shifted to the nearest white matter voxel defined by fractional anisotropy (FA) values of ≥ 0.3. Spheres with 5 mm radius around the selected coordinate served as seed regions. To distinguish dorsal and ventral pathways, coronal slices crossing dorsal tracts at y = 3 to 5 and y = −2 to 0 and crossing ventral tracts at y = 3 to 5 and y = −22 to −19 were manually marked as waypoint masks in MNI space and then morphed into participants’ native space.

Probabilistic tractography between IFG-MTG and PrCG-SPL via dorsal or ventral waypoint masks in each hemisphere was computed bidirectionally using the PROBTRACKX2 module in FSL, with 5000 streamlines per seed region voxel, a curvature threshold of 0.2, step length of 0.5, and maximum number of steps of 2000. Resulting tractography images were cleaned for random connections (threshold at 5% of the image’s maximum intensity value), normalized to MNI space, binarized, and summed. Group level images were slightly smoothed (Gaussian filter with 0.5 mm FWHM), and corrected for filter-induced blurring at the rim (binarization threshold at 0.0001). Plots of pathways found in more than 50% of the participants were generated using brainGL (http://braingl.googlecode.com). Fibre tracts were labelled following the JHU White-Matter-Tractography Atlas in FSL (Hua et al. 2008).

## Supplementary material

**Figure S1.**
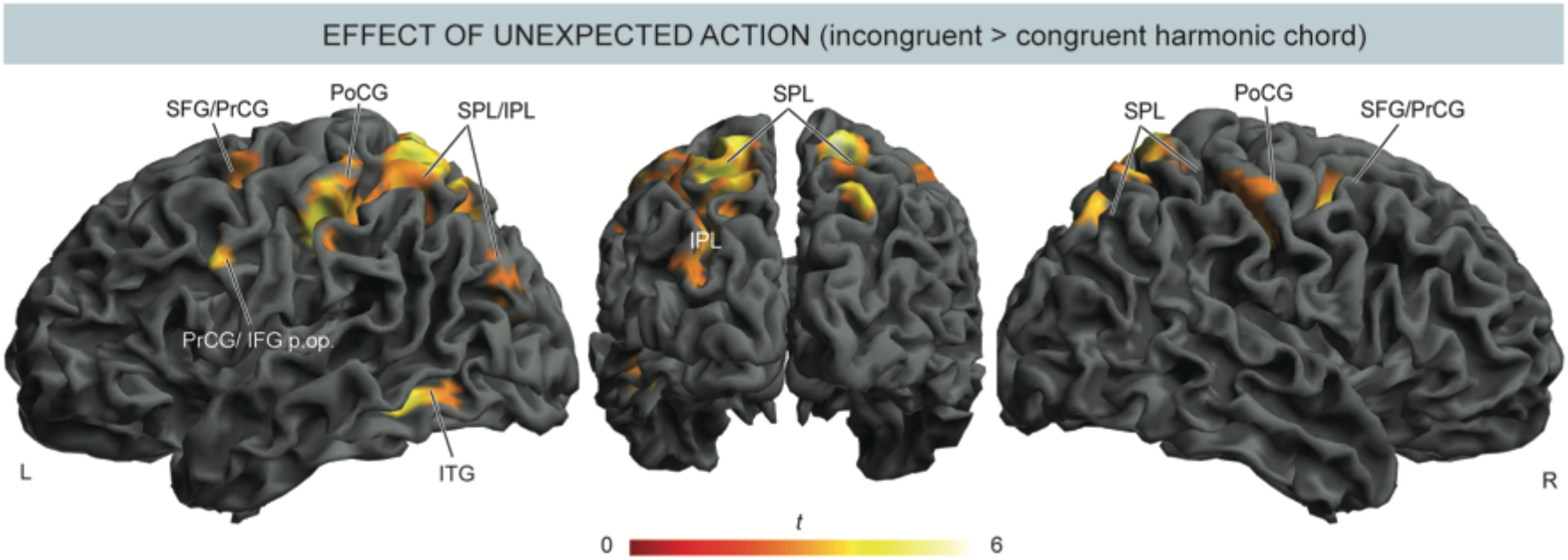
Effect of executing an unexpected action (‘baseline’ blocks). Execution of structurally incongruent chords evoked greater activity than congruent chords in bilateral parieto-frontal action areas. These included left precentral gyrus (PrCG, BA6) and inferior frontal gyrus (IFG, pars opercularis, BA44), bilateral superior frontal gyrus (SFG), postcentral gyrus (PoCG), superior parietal lobule (SPL), and left inferior parietal lobule (IPL) and inferior temporal gyrus (ITG). These findings replicate the results of Bianco et al. (2016) and closely resemble the activity pattern found in the present study following motor violations. This suggests that local structural incongruities also disrupt familiar motor patterns, or put differently, that frequently co-occurring structural elements settle in overlearned motor chunks. BA: Brodmann area.

**Table S1.**
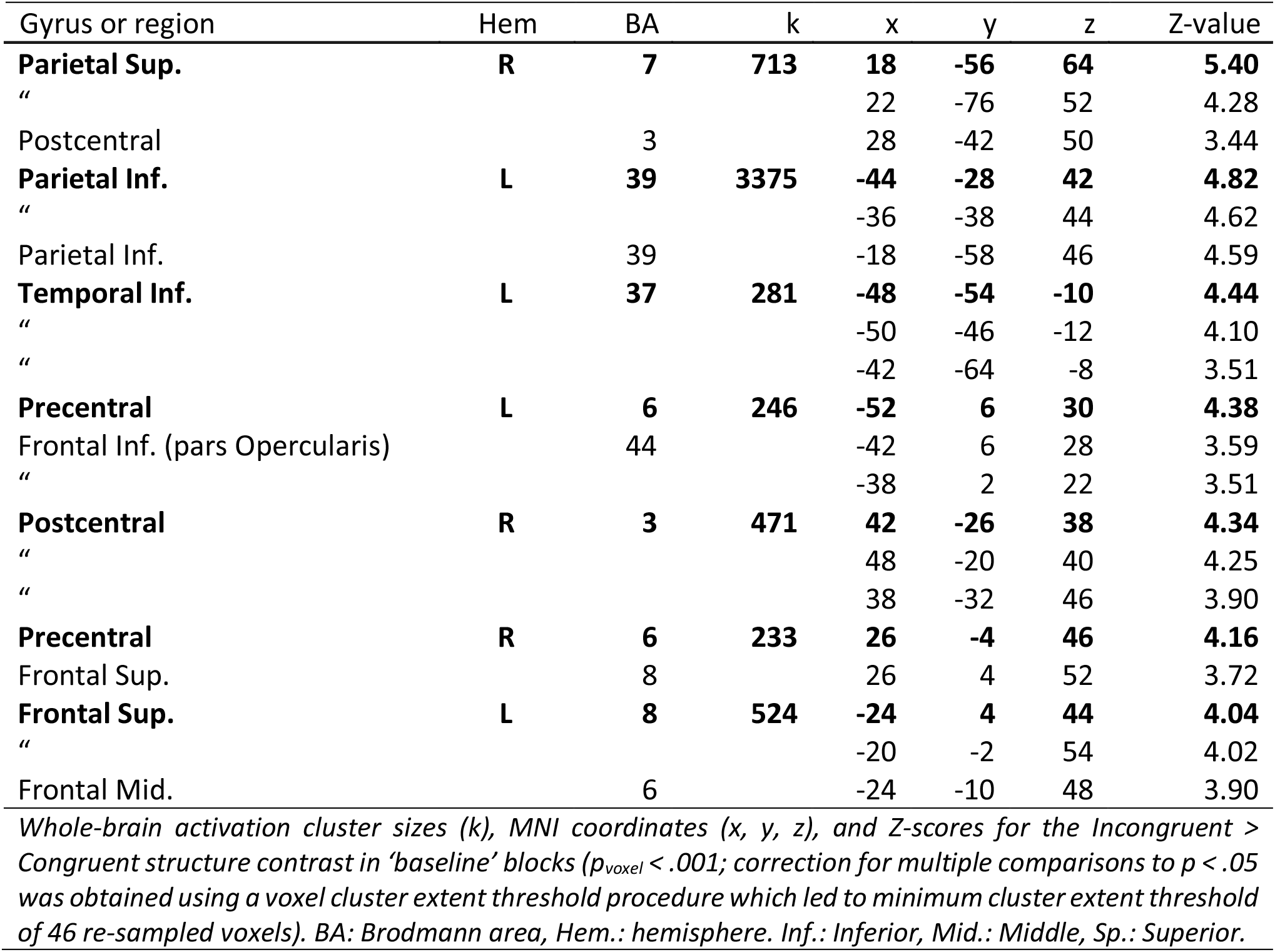
Effect of executing an unexpected action (incongruent > congruent chords).

